# A New Approach for Discovering Functional Links Connecting Non-Coding Regulatory Variants to Gene Targets

**DOI:** 10.1101/2024.06.13.598913

**Authors:** Hammad Farooq, Lin Du, Pourya Delafrouz, Wei Jiang, Constantinos Chronis, Jie Liang

## Abstract

Genome-wide association studies (GWAS) have linked thousands of genetic variants to various complex traits or diseases. However, most identified variants have weak individual effects, are correlated with nearby polymorphisms due to linkage disequilibrium (LD), and are located in non-coding cis-regulatory elements (CREs). These characteristics complicate the assessment of the direct impact of each variant on tissue specific gene expression and phenotype. To address this challenge, we have developed a novel algorithm that leverages polymer folding and 3D chromatin interactions to prioritize and identify putative causal variants and their target genes. From the millions of eQTL-Gene pairs identified by GTEx in human somatic tissues, we classify only ∼10-20% as putative functional eQTL-Gene pairs supported by phenotypic associations confirmed through CRISPR deletion experiments. Our findings show that unlike most variants, functional eQTL-Gene pairs predominantly reside within the same topologically associating domain (TAD) and have strong associations with cell-type specific cis-regulatory elements (CREs), enriched for binding sites of tissue-specific transcription factors. Unlike most approaches that rely on linear distance or other chromatin features (histone code, accessibility), our algorithm emphasizes the importance of physical interactions and 3D chromatin folding in gene regulation, as the identified eQTL-Gene pairs are all among the small fraction of physical chromatin interactions sufficient for chromatin locus folding. Overall, our algorithm reduces false positive associations between DNA variants and genes identified by eQTL analysis and uncovers novel variant-gene pair associations. These findings suggest a mechanism where a small number of regulatory variants control tissue specific gene expression via their physical association with target genes confined within the same TAD. Our approach provides new insights into the molecular mechanisms driving GWAS phenotypes.

## Introduction

Genome-wide association studies (GWAS) reveal associations between thousands of genetic variants and complex traits or diseases [1], [2]. However, only rarely we are able to define the causal variants and their target genes, which hinders the mechanistic understanding of the molecular pathways connecting genetic variants to common diseases [3], [4]. Due to linkage disequilibrium (LD) with nearby polymorphisms, multiple variants can appear to contribute to a phenotype simply owing to a correlation with a causal site [5], [6], [7]. Most of these variants have individually weak effects [8], [9] make it difficult to pinpoint the ‘driver’ or most contributing ones. Statistical fine-mapping [10], [11], [12] can narrow down candidate causal variants but requires a large sample size. Even the largest GWAS studies rarely achieve single-SNP resolution. Moreover, over 90% of these variants are situated within noncoding sequence [13], which may regulate distal genes [14], [15], making it difficult to identify the target causal gene. Perturbation experiments can link variants at cis-regulatory elements (CREs) to target genes but are low throughput [16]. Systematic efforts to pinpoint functional and causative disease-associated variants and their target genes on a larger scale would enhance our understanding of human disease etiology and provide a target for therapeutic interventions.

Extensive efforts by ENCODE [17], Roadmap Epigenomics [18], and others [13], [19] have shown that disease-risk-associated variants are enriched at cis-regulatory elements (CREs) which are only active in relevant cell types, highlighting important regulatory roles for noncoding variants. Inferring target genes of CREs is challenging because CREs can regulate distant genes, making ‘nearest gene mapping’ often ineffective [14], [15], [20], [21]. Recent studies using chromosome conformation capture methods have shown that disease-associated noncoding elements interact with corresponding genes through long-range chromatin interactions [22], with interacting loci enriched for disease-associated SNPs [23], [24]. This suggests that noncoding variants may influence target genes via 3D chromatin interactions, emphasizing the need for high-resolution cis-regulatory interaction maps to understand how these variants affect gene expression.

While eQTL analysis can help to establish genetic connections between regulatory elements (CREs or SNPs) and their respective target genes [25], chromosome conformation methods can establish physical connections between them [26]. Together, these two approaches offer a unique route to mechanistic understanding of how genetic variants affect gene expression by establishing the physical basis of their 3D spatial organization. While important progress has been made in understanding the physical basis of genetic associations between eQTLs and their target genes, existing studies have certain limitations. Studies based on HiChIP [24], [27] and PCHi-C [28] techniques focus solely on the subset of eQTLs that overlap with active cis-regulatory elements or promoter interacting regions, respectively. Consequently, they do not provide information on eQTLs outside of the specific capture regions. On the other hand, Hi-C offers an unbiased view of all possible pairwise interactions between genomic loci genome-wide [29] making it more suitable for generating hypotheses and discovering novel interactions. However, studies employing Hi-C techniques [30], [31] have typically remained at the levels of topologically associating domains (TADs) rather than extensively exploring the chromatin interactions involving eQTLs. Furthermore, quantifying chromatin interactions involving eQTLs using Hi-C is challenging due to the potential presence of numerous chromatin contacts, many of which may arise from generic effects of confined polymers [32], [33]. Existing methods integrating Hi-C with eQTL data are also unable to effectively remove these random polymer effects.

In this study, we developed a novel computational pipeline to link genetic variants with their target genes by integrating genetic and physical interaction data. Our approach prioritizes putative functional variants and genes that exhibit both genetic and physical interactions. Our findings show that unlike most variants, functional eQTL-Gene pairs predominantly reside within the same topologically associating domain (TAD) and have strong associations with cell-type specific cis-regulatory elements (CREs), enriched for binding sites of tissue-specific transcription factors. Furthermore, these prioritized variants show higher enrichment in causal variants identified by reporter assays, statistically fine-mapped eQTLs, and disease-associated GWAS. They are also enriched with motifs for tissue-specific transcription factors, which in turn are linked to genes involved in tissue-specific biological processes. CRISPR-based experimental data further validates the variant-gene map predicted by our method. Our physical proximity-based approach predicts the long-range target genes of enhancers more accurately than state-of-the-art methods.

Overall, our study provides a physical basis for genetic associations by systematically linking genetic variants with their target genes. This enables the identification of functional regulatory variants and offers a mechanistic understanding of how genetic variants affect specific tissues or gene expression.

## Results

### A Small Proportion of eQTLs Are Identified as Physical eQTLs

GTEx analysis has revealed a significant number of eQTL-eGene associations, ranging from 0.4 to 2.4 million across different tissues (Supplementary Fig. S1c). On average, this translates to over 100 eQTLs per eGene. Identifying functionally important eQTLs from this vast pool presents a formidable challenge. Deciphering the functional relevance of these associations is therefore inherently complex. Chromatin interactions enable the spatial juxtaposition of distal cis-regulatory elements and promoters, they can directly modulate the expression of the target gene [22]. We hypothesized that eQTLs exhibiting both genetic and physical interactions with their target genes may have enhanced functional significance.

Assessing whether physical interactions exist between two genomic loci through Hi-C data poses significant challenges. Hi-C studies often record numerous chromatin contacts, but majority of which may arise from generic effects of confined polymers [32], [33]. To identify Hi-C interactions unlikely to occur due to such random collisions, we employed the CHROMATIX method [33]. We begin by generating an ensemble of random self-avoiding chromatin polymers under nuclear confinement using the fractal Monte Carlo sampling method [33]. We then bootstrap over this ensemble, which serves as our null model of random physical interaction, to assign *p-values* to Hi-C interactions. Those interactions with BH-FDR adjusted *p-values* below a threshold of 0.05 are considered significant (see Fig. 1b). These significant non-random interactions have been demonstrated to be able to drive chromatin folding (see Supplementary Fig. S1b) [33], [34], [35]. In addition they have been shown to enable the identification of specific chromatin many-body interactions [33] and the quantification of heterogeneity in 3D chromatin structures within the cell population [35]. We refer these significant non-random interaction as folding reconstitutive (FoldRec) interactions.

**Figure 1:**
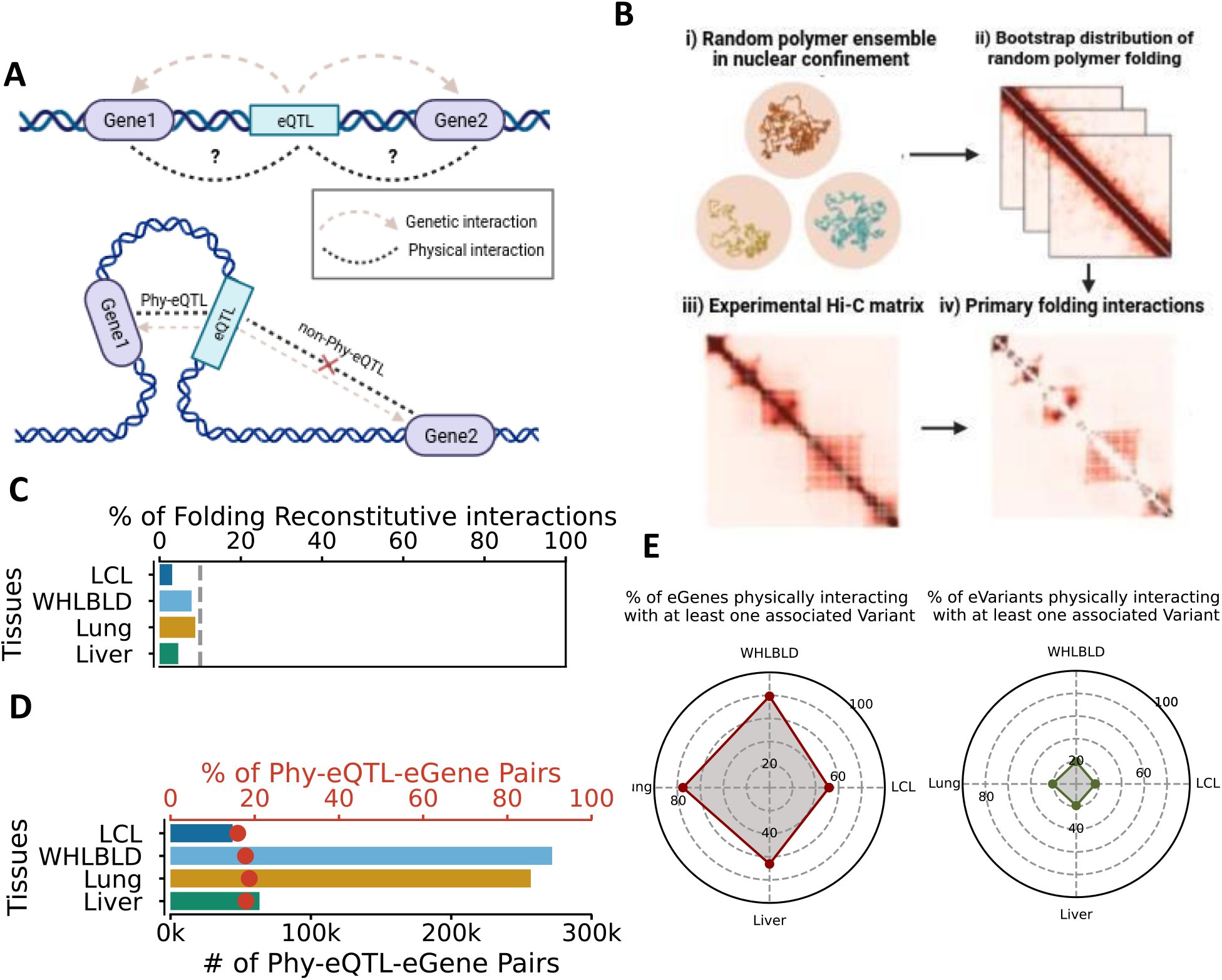
Small Percentage of eQTL identified as Phy-eQTLs. **(a)** Schematic representation distinguishing Phy-eQTL and non-Phy-eQTL. Phy-eQTL involves both genetic and physical interactions with the target gene (eGene1), while non-Phy-eQTL lacks physical interaction despite its genetic association with the gene (eGene2). **(b)** Steps involved in identifying FoldRec interactions from Hi-C data. It includes generating a random polymer ensemble, creating a null model with a bootstrap distribution, and filtering out background interactions to identify FoldRec interactions. **(c)** Percentages of the Hi-C interactions identified as FoldRec interactions. **(d)** Percentage and number of the eQTL-eGene pairs identified as Phy-eQTL-eGene. The top axis represents the percentage, while the bottom axis represents the count of Phy-eQTL-eGene pairs. **(e)** Left: Percentage of eGenes physically interacting with at least one eVariant. Right: Percentage of eQTL physically interacting with at least one eGene.

Here, we carry out our study on four GTEx [36] tissues, EBV-transformed lymphocytes (LCL), whole blood (WHLBLD), lung, and liver with corresponding 4DN Hi-C datasets of GM12878, K562, IMR90, and HepG2 cell lines, respectively (Supplementary Fig. S1a) [37], [38]. These tissues were selected due to the availability of both Hi-C and eQTL datasets, allowing us to apply our approach across a variety of tissues to ensure robustness and comprehensiveness. Across all tissues, only a minimal proportion, ranging from 3% to 9% of Hi-C interactions were identified as FoldRec interactions (See Fig. 1c), consistent with the earlier studies [33], [35].

We investigate whether an eQTL and its associated eGene exhibit a 3D physical interaction, we categorize eQTL-eGene pairs based on whether FoldRec interaction exists between them or not. If both genetic and FoldRec interactions exist between an eQTL and its associated eGene, we referred to them as <u>phy</u>sical-<u>eQTL</u> (PhyeQTL) and physical-eGene (PhyeGene), respectively. Conversely, if only genetic association exists, we referred them as non-<u>phy</u>sical-<u>eQTL</u> (non-PhyeQTL) and non-physical-eGene (non-PhyeGene), respectively (refer to Fig. 1a).

Among all eQTL associations identified by GTEx, approximately 11% to 16% correspond to pairs where the eQTL is positioned within 10kb from the transcription start site (TSS) of the associated eGene (Supplementary Fig. S1d). We focus on long-range interaction and exclude them from our analysis due to the resolution limitations of Hi-C data. We overlay the long range (>10kb) eQTL-eGene pairs with FoldRec interactions to determine whether the eQTL physically interacts with the promoter of the eGene. Our results show that about 52% to 79% of eGenes harbor at least one PhyeQTL, and 17% to 21% of eQTLs are associated with at least one PhyeGene (Fig. 1e). However, only a small percentage of all eQTL-eGene pairs, ranging from 16% to 19%, are identified as PhyeQTL-Gene (Fig. 1d). These results reveal that only a small fraction of eQTLs are PhyeQTL.

To evaluate the functional significance of those PhyeQTL, we assessed their enrichment in functionally validated genetic variants by Massively Parallel Reporter Assays (MPRAs), specifically SuRE, which systematically screen millions of human SNPs for potential effects on regulatory activity [39]. Our analysis revealed that PhyeQTLs exhibit higher enrichment in reporter assay QTL (raQTL) compared to non-PhyeQTLs, indicating their functional relevance (Supplementary Fig. S2b). This suggests that PhyeQTL can serve as a physically informed approach for fine mapping of eQTLs.

Additionally, we examined whether PhyeQTL are enriched for statistically fine-mapped putative causal variants for expression quantitative loci (eQTL). These variants are identified by the three different methods: CaVEMaN [10], CAVIAR [11] and dap-g [12]. Strikingly, PhyeQTL displayed greater enrichment in causal variants identified by all three methods compared to non-PhyeQTL (Supplementary Fig. S2a), further highlighting the importance of PhyeQTL.

Furthermore, we also assessed whether PhyeQTL exhibits any preference for disease or trait associated GWAS variants. Again, we observed that this enrichment was notably higher for PhyeQTL compared to non-PhyeQTL (Supplementary Fig. S2c).

In summary, our findings demonstrate that, among all eQTL variants, only a small fraction are Physical QTLs. This underscores the potential of utilizing physical interaction information to prioritize causal eQTL variants and GWAS variants.

### Physical eQTL-eGene Pairs Predominantly Fall Within the Same TAD

Several studies have suggested that Topologically Associating Domains (TADs) play a dual role in facilitating and constraining enhancer-promoter interactions [40], [41]. Deletions or rearrangements of TAD boundaries can disrupt the regulation of gene expression by losing or hijacking of regulatory elements [42], [43]. To investigate the relationship between eQTL-eGene pairs and TADs, we stratified eQTL-eGene pairs based on whether they crossed TAD boundaries or remained within the same TAD (Fig. 2a). Pairs where the eQTL and its associated eGene resided within the same TAD were classified as TAD boundary non-crossing, while pairs where they were located in different TADs, thus associated by crossing TAD boundaries, were classified as TAD boundary crossing. We found, over 85% of PhyeQTL-eGene pairs were contained within the same TAD, and a small fraction of ∼15% are crossing the TAD boundary. Conversely, approximately 50% of the non-PhyeQTL-eGene pairs fell into both TAD boundary crossing and non-crossing categories (Fig. 2b). This striking difference between PhyeQTL and non-PhyeQTL suggests that they employ different mechanisms to regulate eGene expression. As, PhyeQTL-eGene pairs predominantly fall within the same TAD, consistent with the observation that enhancer-promoter (E-P) communication occurs primarily within insulated neighborhoods [41], [43]. These results suggest the direct effects of PhyeQTL on eGene expression. In contrast, non-PhyeQTL exhibit no relationship with TADS, suggesting that they may act through more indirect mechanisms to influence eGene expression.

**Figure 2:**
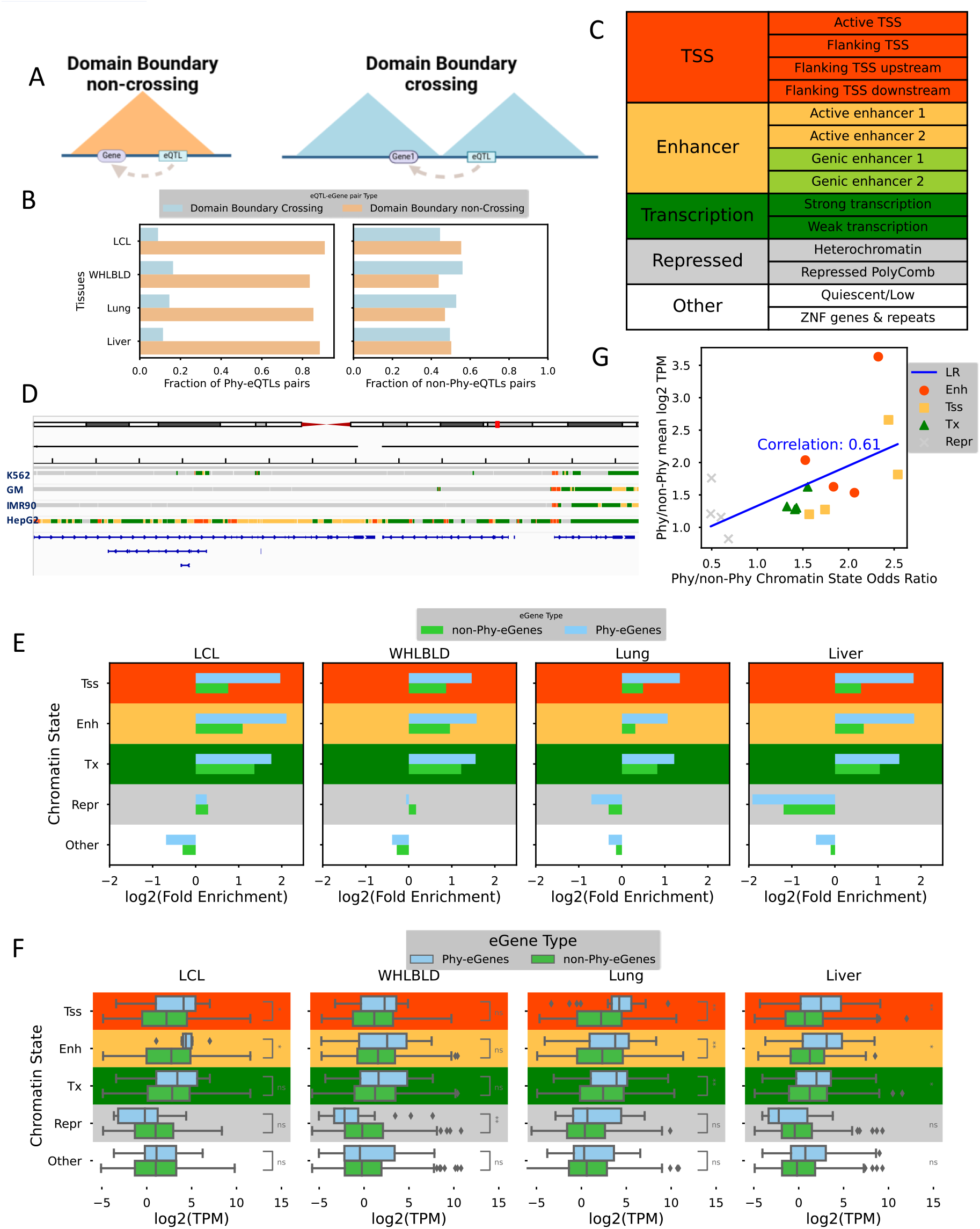
Enhanced Gene Expression Levels of Phy-eGenes are Linked to Active Chromatin Enrichment of Phy-eQTLs. **(a)** Left: Illustration of a Domain boundary non-crossing scenario where the pair of eQTL-eGene lies within the same Hi-C domain. Right: Illustration of a Domain boundary crossing scenario where the pair of eQTL and its associated eGene are situated in different Hi-C domains. **(b)** Proportion of eQTL-eGene pairs in both categories: Domain boundary crossing and Domain boundary non-crossing. Left: Phy-eQTL pairs. Right: non-Phy-eQTL pairs. **(c)** Representative chromatin states, summarized by merging similar states of 15-state CHROMHMM state model. **(d)** Chromatin states annotation of an example locus. **(e)** Log-fold enrichment of Phy-eQTL and non-Phy-eVariant over all eQTL genotyped in GTEx, categorized by the 5 chromatin states. Highlighting the active chromatin enrichment of Phy-eQTLs. **(f)** Gene expression comparison between Phy-eGenes and non-Phy-eGenes, stratified by the chromatin state annotation of the associated eVariant. Highlighting the enhanced gene expression levels of Phy-eGenes associated with active Phy-eGenes. **(g)** Correlation between chromatin state enrichment and expression of Phy-eQTL/non-Phy-eQTL and Phy-eGene/non-eGene, respectively.

### Enhanced Expression of Target eGenes Linked to Enrichment of Active Chromatin in Physical eQTLs

Previous studies suggest that intergenic eQTLs demonstrate a remarkable tendency to overlap with transcription factor binding sites, open chromatin regions, promoters, and enhancers [44], [45]. To delve deeper into the specifics of the chromatin landscape underlying phy-eQTLs and non-PhyeQTLs. We harnessed the ChromHMM 15-state model [19] form ENCODE [46], consolidating similar states to derive four representative chromatin states (see Fig. 2c). These tissue-specific summarized chromatin state annotations were then utilized to annotate the entire genome, providing insights into tissue-specific chromatin activity (as depicted in Fig. 2d).

Using all variants genotyped in GTEx as a comparative background, we evaluated the enrichment for overlap of PhyeQTL and non-PhyeQTL across each chromatin state. Notably, PhyeQTL and non-PhyeQTL exhibited strikingly divergent levels of enrichment. Specifically, PhyeQTL demonstrated higher enrichment within active chromatin states (TSS, Enh, and Tx), whereas non-PhyeQTL showed greater enrichment within inactive chromatin states (Repressed) (see Fig. 2e). The pronounced differences in the enrichment of PhyeQTL within active regulatory regions highlight the enhanced regulatory potential of PhyeQTL in contrast to non-PhyeQTL. This emphasizes that PhyeQTL SNPs are more inclined to disturb active regulatory regions, thereby amplifying their functional importance.

The noticeable differences in chromatin state enrichment linked with PhyeQTL motivate us to explore whether this increased regulatory capacity correlates with observable impacts on the expression levels of the associated eGenes. To address this question, we conducted an analysis of eGene expression levels, stratified by the chromatin state of the corresponding PhyeQTL or non-PhyeQTL, by aligning the active or inactive chromatin state of eGenes with their respective associated PhyeQTL or non-PhyeQTL. Notably, eGenes associated with active (TSS, Enh and Tx) PhyeQTL display significantly higher expression levels than those associated with active non-PhyeQTL. Conversely, eGenes associated with inactive (Repressed) PhyeQTL exhibit notably lower expression levels compared to eGenes linked to inactive non-PhyeQTL (see Fig. 2f). Moreover, there is a positive correlation between chromatin state enrichment and expression levels of PhyeQTL/non-PhyeQTL and PhyeGene/noneGene, respectively (see Fig. 2g). Overall, these findings suggest that the chromatin state of PhyeQTL directly influences the expression levels of associated eGenes, while non-PhyeQTL do not exert a strong direct influence on expression, implying that PhyeQTL have a greater potential to directly modulate eGene expression compared to non-PhyeQTL. Non-PhyeQTL may instead influence expression through indirect mechanisms. This underscores the significance of PhyeQTL information in elucidating functional connections between regulatory elements and genes within the genome.

### Physical QTLs Likely Disrupt Potential Regulatory Elements

The increased enrichment of PhyeQTL in active chromatin states (Fig. 2e) led us to consider whether PhyeQTL possesses an enhanced ability to impact regions characterized by open chromatin and active enhancers. To investigate this, we assessed whether PhyeQTL sites exhibit greater enrichment for ATC-seq and H3k27ac signals, which are indicators of open chromatin and active enhancers, respectively. Analysis of the average profiles of ATAC signals (Fig. 3a) and H3k27ac signals (Fig. 3b) revealed that PhyeQTLs exhibit higher enrichment of both signals across tissues compared to non-PhyeQTLs. These differences in ATAC and H3k27ac signals imply that PhyeQTLs are proximal to active enhancers and have an increased potential to perturb active enhancer in these regions.

**Figure 3:**
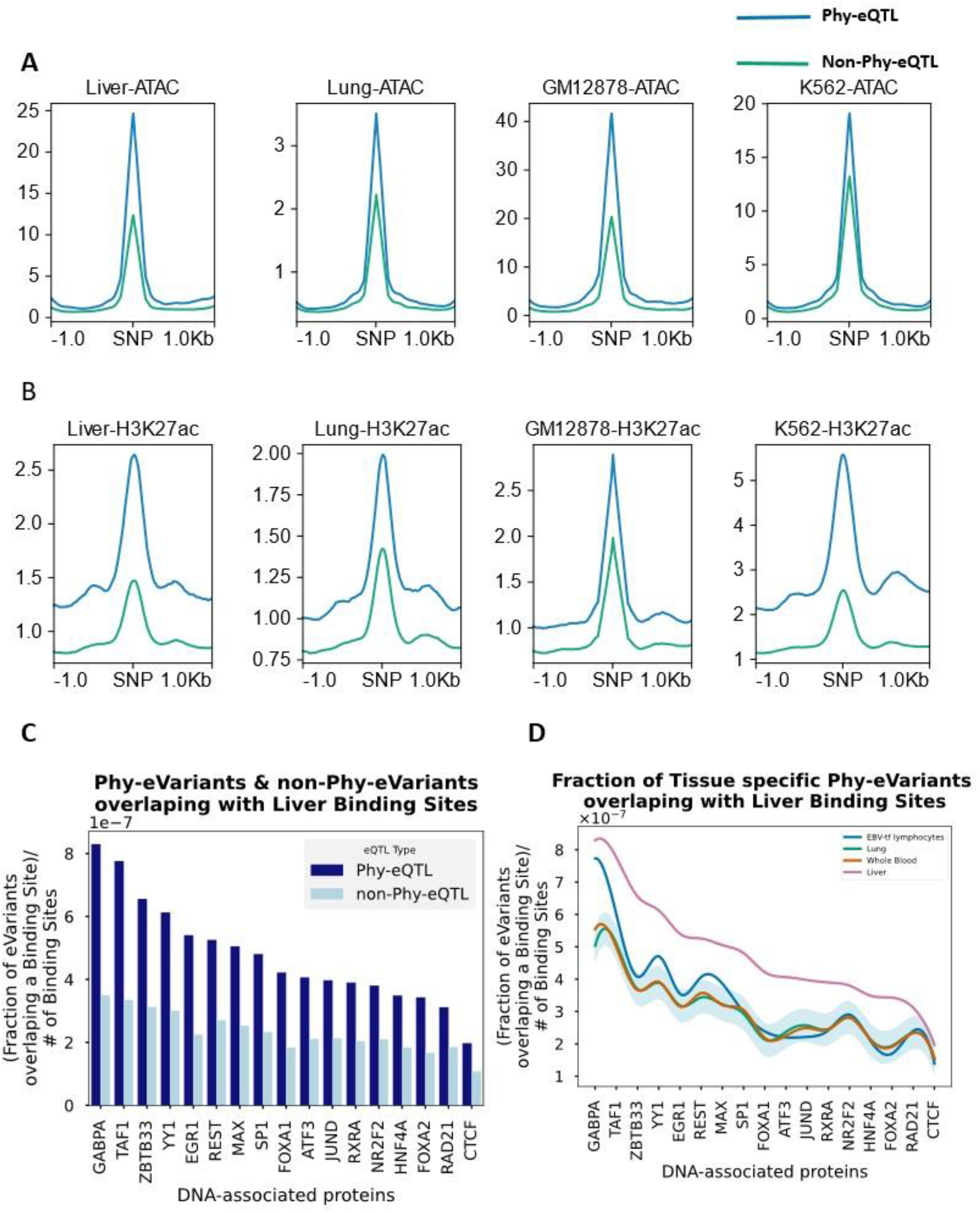
Phy-eQTLs Likely Disrupt Potential Regulatory Elements. **(a)** Average profiles of ATAC signals centered on either eQTLs or peak summits, when within a 1kb proximity to the eQTL, highlighting the potential disruption of accessibility by Phy-eQTLs. **(b)** Average profiles of H3k27ac signals centered on either eQTLs or peak summits, when within a 1kb proximity to the eQTL, highlighting the potential disruption of active enhancers by Phy-eQTLs. **(c)** Fraction of liver eQTLs overlapping with the binding sites of the DNA-associated protein, normalized by the total number of binding sites. Liver Phy-eQTLs exhibit a predominant overlap with binding sites compared to non-Phy-eQTLs, highlighting the potential disruption of binding sites of DNA-associated protein by Phy-eQTLs. **(d)** Normalized fraction of Phy-eQTLs for different tissues overlapping with DNA-associated protein binding sites of the Liver tissues. Liver Phy-eQTLs exhibit a predominant overlap with binding sites compared to other tissues, suggesting a potential disruption of tissue-specific binding sites by Phy-eQTLs.

Additionally, we examined whether PhyeQTLs affect the binding sites of DNA-associated proteins (DAPs), such as transcription factors (TFs), which bind to genomic regulatory elements such as promoters, enhancers, silencers, and insulators [47]. We utilized ChIP-seq data obtained from liver tissue to quantify the binding of 17 different DAPs [48], including liver-specific factors (e.g., HNF4A and RXRA) and chromatin structure-related factors (e.g., CTCF and RAD21). Our analysis aimed to evaluate the overlap between QTLs and DAP binding sites, and comparing the overlap patterns between PhyeQTLs and non-PhyeQTLs. Specifically, we calculated the fraction of liver eQTLs overlapping with the binding sites of DAPs, normalized by the total number of binding sites for each DAP. Remarkably, Liver PhyeQTLs exhibited a predominant overlap with binding sites compared to non-PhyeQTLs. This strongly suggests that PhyeQTLs can disrupt the DAP binding sites (Fig. 3c and Fig. S3).

We further ask whether the disruption of DAP binding sites by PhyeQTLs is tissue specific. As reported by Ramaker *et al*., the data on DAP binding sites derived from liver tissue closely reflects liver biology than that from the HepG2 cell line [48]. Given the unavailability of primary tissue-based data on DAP binding sites for other tissues analyzed in this study, we utilized liver tissue-derived DAP binding site data to assess tissue-specificity and evaluate the overlap enrichment of PhyeQTLs from other tissues with DAP binding sites. Notably, Liver PhyeQTLs exhibited a prominent overlap with binding sites of Liver DAPs compared to PhyeQTLs from other tissues (see Fig. 3d), suggesting a potential disruption of tissue-specific binding sites by PhyeQTLs.

In summary, these results reinforce the idea that PhyeQTLs exhibit increased regulatory capacity and are prone to disturb active regulatory elements, such as open chromatin, active enhancers, and tissue-specific TF binding sites.

### Physical eQTLs are Enriched with Tissue-specific Motifs

The enrichment of PhyeQTL overlapping with liver DAP binding sites underscores the importance of identifying all putative transcription factors (TFs) binding at these sites to understand how PhyeQTL disrupt regulatory mechanisms. However, the lack of comprehensive tissue-specific DAP binding data across all four tissues analyzed in this study presents a formidable challenge in identifying these TFs. To overcome this challenge, we aim to quantify the enriched TF motifs at PhyeQTL sites, thereby identifying putative TFs bound to these regions.

We employed the HOMER algorithm [49] for de novo motif discovery to uncover DNA motifs overrepresented within PhyeQTL sites. In each tissue, we conducted motif enrichment analysis on all sequences within PhyeQTL sites (± 100 bp), taking sequences from non-PhyeQTL sites (± 100 bp) identified within that tissue as background. This enables the identification of motifs enriched specifically at PhyeQTL sites compared to non-PhyeQTL sites, facilitating insight into their regulatory significance.

For each identified TF motif, we retained those TFs expressed in the corresponding GTEx tissue (median TPM > 1). The collective analysis of TF motifs identified across all tissues underscores their tissue specificity both in motif enrichment and expression (see Supplementary Fig. S4 and Fig. S5). Notably, in LCL, we detected motifs corresponding to transcription factors implicated in B-cell development and immune responses [50], including PAX5 [51], NF-KB [52] and SP1/PU.1 [53]. In Whole Blood, we identified motifs associated with immune and inflammatory responses, as well as hematopoiesis and erythroid development, such ETS-1, ETS-2 [54], NF-KB1 [55] and GATA-1 [56]. In lung tissue, motifs linked to lung development and morphogenesis emerged, including GATA6 [57], FOXA2 [58] and CEBPB [59]. Lastly, in liver tissue, motifs corresponding to liver development-associated transcription factors [60], [61] were identified, including HNF4A, CEBpB, and FOXA3.

Motif enrichment analysis in each tissue unveiled that PhyeQTL sites exhibit enrichment with binding sites of tissue-specific TFs. This observation prompted us to delve deeper into the PhyeGenes linked to PhyeQTL that coincide with TF binding sites. To systematically conduct this analysis, we selected the top 10 expressed TFs (median TPM) from the pool of all TF motifs identified within a tissue (Fig. 4a). Subsequently, we scanned for instances of these motifs overlapping with PhyeQTL and retrieved the corresponding PhyeGenes. Taking these PhyeGenes as a gene set, we performed gene list enrichment analysis using Enrichr [62]. Our analysis revealed that these PhyeGenes are involved in tissue-specific GO Biological Processes (Fig. 4b). Specifically, GO terms associated with LCL and whole blood included immune response and antigen presentation, among others. Liver-related GO terms included blood fluid secretion and metabolic processes. Conversely, lung-related GO terms exhibited relatively less tissue specificity. Overall, these findings underscore the functional significance of PhyeQTL, as they are enriched with motifs for tissue-specific TFs, which in turn are linked to PhyeGenes involved in tissue-specific biological processes.

**Figure 4:**
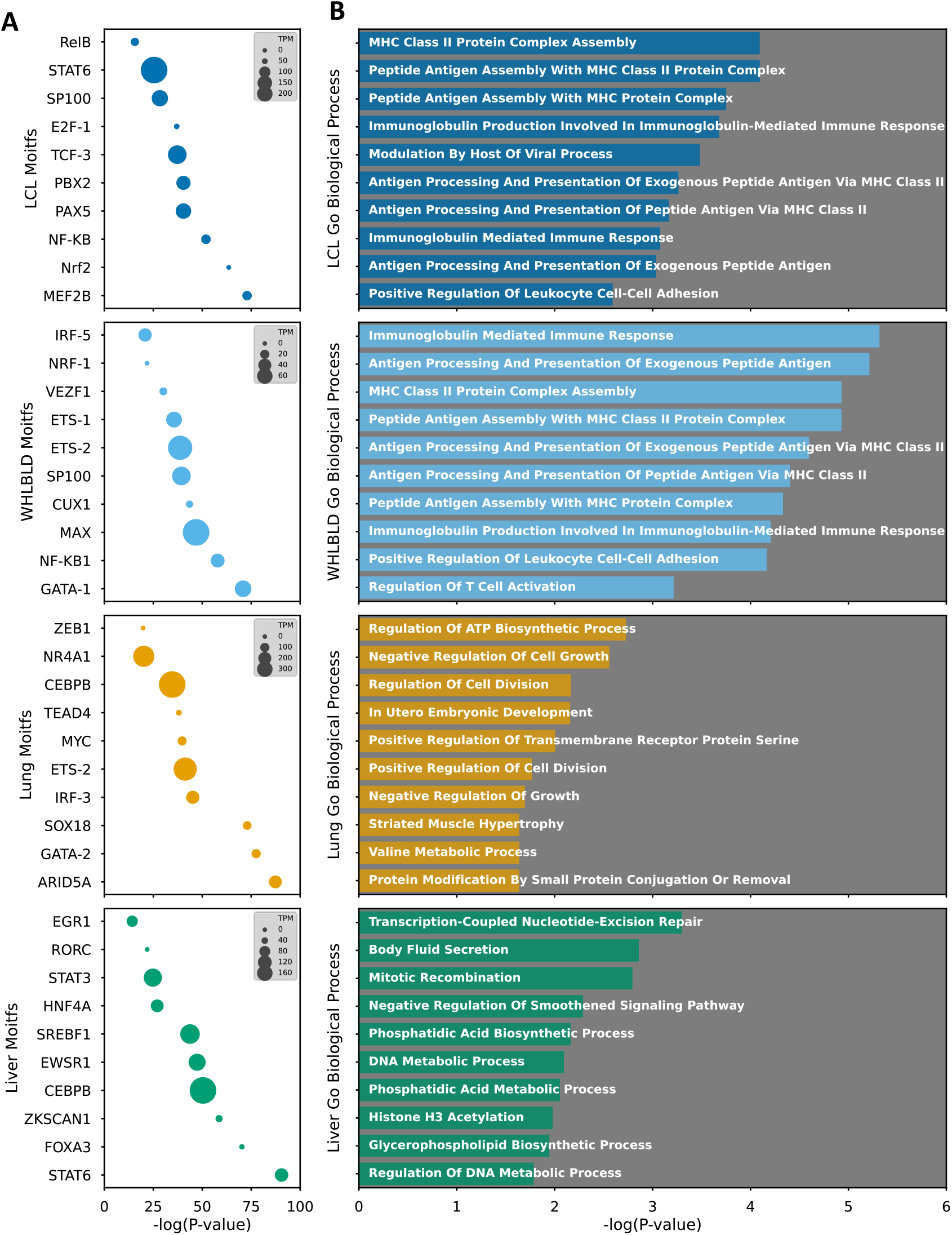
Phy-eQTLs are enriched with Tissue-specific Motifs. **(a)** Motif enrichment was computed on the Phy-eQTL sites for each tissue. From all identified motifs, only the top 10 expressed transcription factors (TFs) in the corresponding tissue are plotted. The x-axis represents the enrichment (-log(p-value)) computed by HOMMER, while the size of the scatter points represents the expression (TPM) of the corresponding motif. **(b)** Gene Ontology (GO) enrichments of a subset of Phy-eGenes associated with Phy-eQTLs containing one of the top ten tissue-specific motifs depicted in (a) for each tissue.

In summary, our results suggest that PhyeQTL influences the expression of tissue-specific PhyeGenes by disrupting the binding sites of tissue-specific TFs.

### Tissue-specific Enh Phy-eQTL modulates tissue-specific gene expression

As PhyeQTLs exhibit characteristics typical of active cis-regulatory elements (CREs), including enriched active chromatin states, open chromatin, and H3k27ac signals. Moreover, they significantly overlap with tissue-specific DNA binding sites for DAP (DNA-associated proteins) and transcription factor motifs known to regulate tissue-specific genes. These distinct features of PhyeQTLs have prompted us to leverage their potential for constructing tissue-specific enhancer-gene maps.

Our approach begins by annotating tissue-specific PhyeQTLs within the respective tissue as well as across other tissues. Remarkably, over 75% of Enhancer (Enh) PhyeQTLs are exclusively annotated as active enhancers in a single tissue, underscoring the tissue-specific regulatory role of PhyeQTLs (Fig. 5a). Furthermore, the PhyeGenes associated with tissue-specific Enh PhyeQTLs in each tissue exhibit minimal overlap with those in other tissues, reinforcing the tissue-specific nature of their regulatory activity (Fig. 5b). This observation prompted us to investigate the expression patterns of PhyeGenes across various tissues.

**Figure 5:**
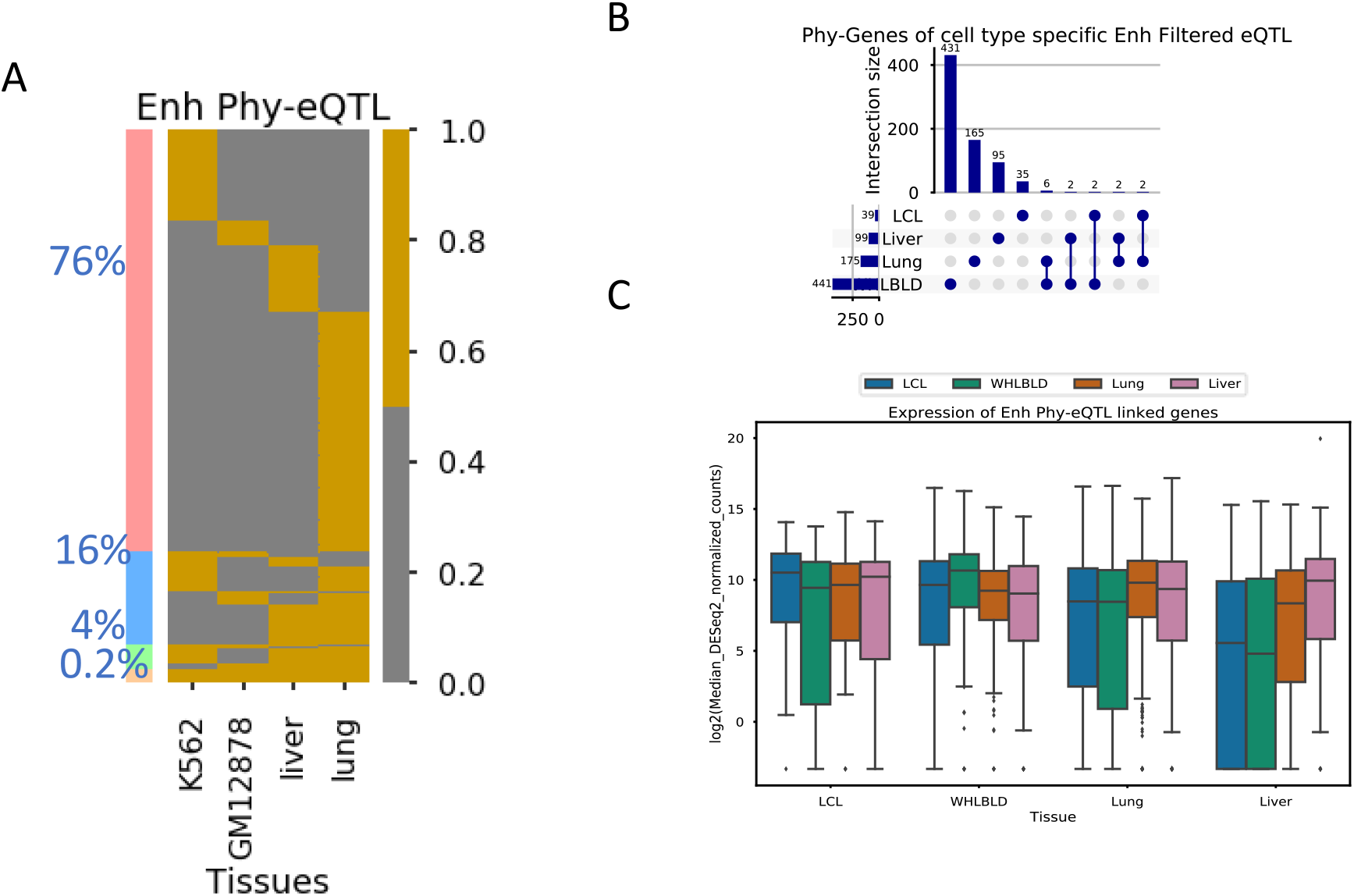
Tissue-specific Enh Phy-eQTL modulates tissue-specific gene expression. **(a)** Percentage of Phy-eQTL annotated as Enhancer state across one tissue (76%), two tissues (16%), three tissues (4%), and all four tissues (0.2%). **(B)** UpSet plots illustrating the intersection of Phy-eGenes associated with Tissue-specific Enh Phy-eQTL. **(C)** Comparative gene expression analysis between Phy-eGenes associated with Tissue-specific Enhancer Phy-eQTL and the corresponding genes in other tissues. Highlighting the enhanced gene expression levels of Phy-eGenes linked to Tissue-specific Enhancer Phy-eQTL in contrast to those observed in other tissues.

Comparative gene expression analysis between PhyeGenes associated with tissue-specific Enhancer PhyeQTLs and their counterparts in other tissues shows significantly higher expression levels of PhyeGenes linked to tissue-specific Enhancer PhyeQTLs, compared to those in other tissues (Fig. 5c). These findings collectively underscore the potential of PhyeQTLs in identifying cis-regulatory elements and their target genes.

### Validation of Physical eQTLs-eGene Associations Using CRISPR Data

For functional validation of our PhyeQTL-derived enhancer-gene maps, we utilized CRISPR-based experimental data from K562 cells [16] and integrated it with our PhyeQTL-PhyeGene pairs identified in whole blood. Our analysis, employing FoldRec, identified MYO1D and TMEM98 as PhyeGenes associated with PhyeQTLs. Notably, these PhyeQTLs colocalized within the same 5kb Hi-C bin as experimentally validated enhancer elements for MYO1D and TMEM98 genes (Fig. 6a), thereby validating the accuracy of our PhyeQTL-based enhancer-gene maps.

**Figure 6:**
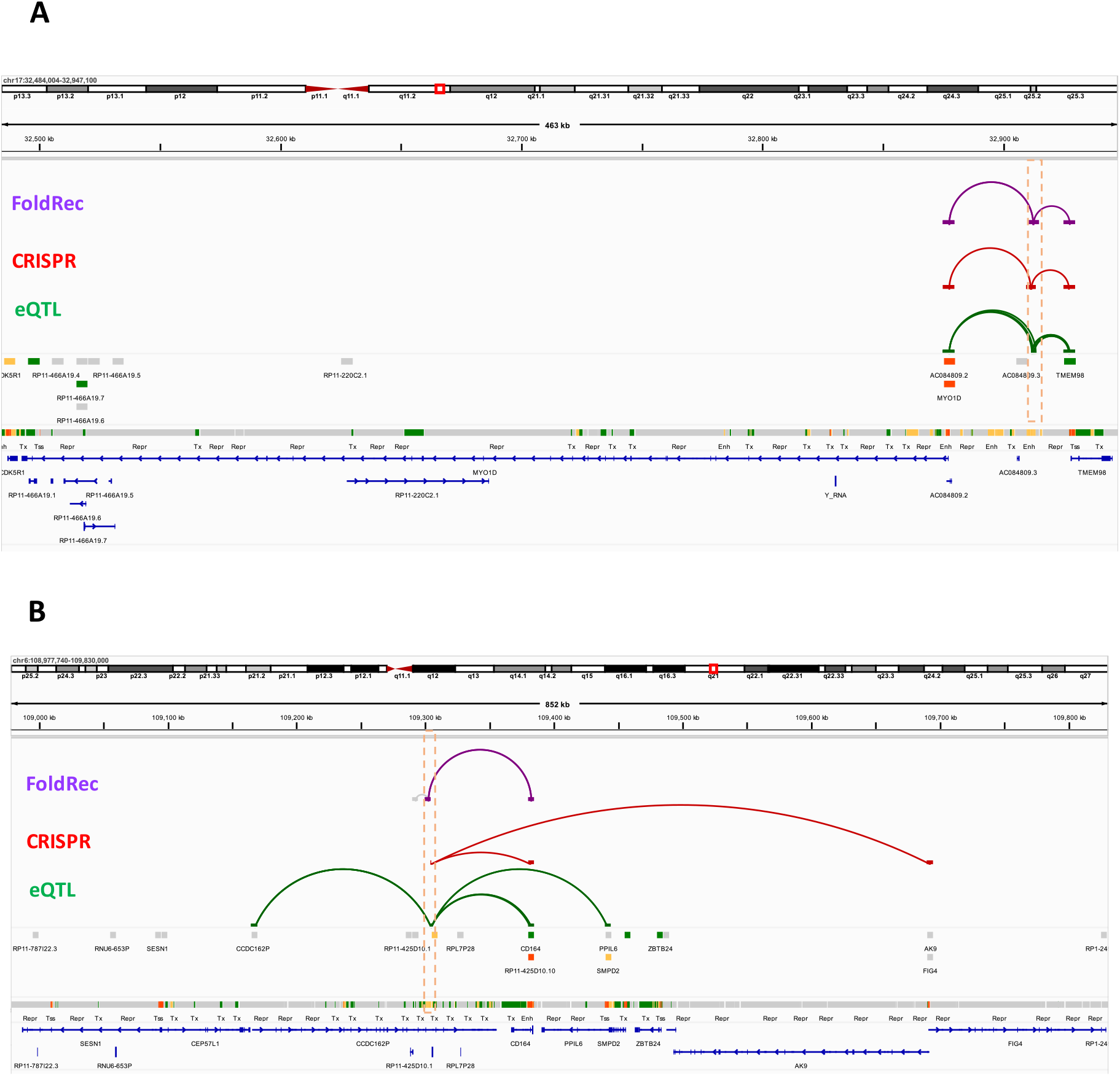
Phy-eQTL-gene association validated by Crispr data. **(a)** A set of Phy-eQTLs, co-located within the same Hi-C bin, exhibiting physical interactions with eGenes MYO1D and TMEM98, demonstrating consensus with Crispr data. **(b)** A set of Phy-eQTLs, co-located within the same Hi-C bin, exhibiting physical interactions with eGene CD164 demonstrating consensus with Crispr data.

Similarly, we identified CD164 as a PhyeGene associated with PhyeQTL. This PhyeQTL was found to co-locate with a CRISPR-validated enhancer element of the CD164 gene within the same Hi-C bin (Fig. 6b), further validating the reliability of our PhyeQTL-derived enhancer-gene maps.

These instances of functional validation highlight the robustness of PhyeQTLs in identifying functional enhancer-gene associations, thereby emphasizing their potential in uncovering the regulatory landscape of the genome.

### Identification of Targets of eQTLs Beyond Their eGenes Validated by CRISPR Data

A significant portion, approximately 80%, of eQTLs do not exhibit direct physical interactions with any of their associated eGenes (Fig. 1d). To gain deeper insights into how non-PhyeQTL exert their influence on the expression of target eGenes, we seek to identify the potential alternative interacting partners of these non-PhyeQTL when they do not directly interact with the associated eGene (Fig. 7a). These partners referred here as “other integrating end of non-PhyeQTL”. Using whole genome as a comparative background, we evaluated the chromatin state enrichment of these other integrating end of non-PhyeQTL. Our analysis reveals a positive log-fold enrichment of active chromatin states (TSS, Enh, and Tx) in these regions, indicative of their regulatory potential.

**Figure 7:**
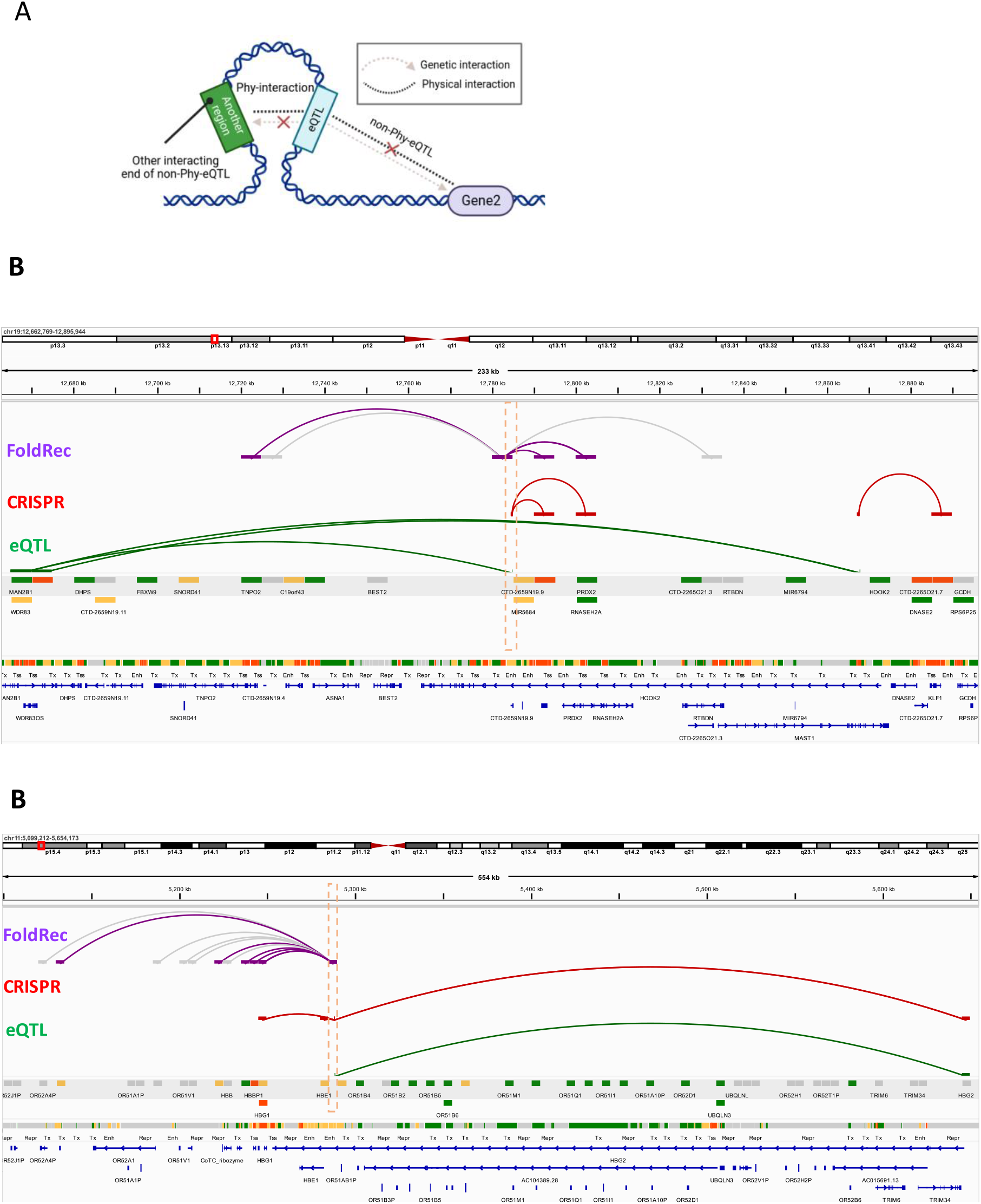
FoldRec Identified Targets of Non-Phy-eQTLs Beyond Their eGene which are Validated by Crispr Data. **(a)** Schematic representation of the other interacting end of the non-Phy-eQTLs. These are the genomic regions (other than genetically associated eGene) that physically interacting with the non-Phy-eVariant. **(b)** A set of two non-Phy-eQTLs, collocated within the same Hi-C bin, demonstrating physical interactions with JUNB and PRDX2, which are non-eGenes of these eQTLs. These interactions are validated by Crispr data. **(c)** A non-Phy-eQTLs demonstrating physical interactions with HBG1, which is non-eGene of this eQTL. This interaction is validated by Crispr data.

Based on these findings, we propose a hypothesis that non-PhyeQTL may exert regulatory effects on genes beyond those directly associated with eQTL. To test this hypothesis, we integrated CRISPR-based experimental data from K562 cells [16] in our analysis of alternative interacting partners of non-PhyeQTL.

Remarkably, functional enhancers of JUNB and PRDX2 identified by CRISPR are within the same Hi-C bin as two non-PhyeQTLs, namely chr19_12784786_C_G_b38 and chr19_12784786_C_G_b38 (Fig. 7b). Although these eQTLs are not genetically associated with JUNB and PRDX2, the physical interaction with these genes suggests their functional relevance. Similarly, we observed a non-Phy-eQTL interacting physically with HBG1, a gene not associated with this eQTL. This finding is also supported by CRISPR data (Fig. 7c). These functional data validate our hypothesis that non-PhyeQTL influences the expression of non-eGenes through physical interaction.

Our findings suggest two possibilities: non-PhyeQTL may indirectly impact eGenes through alternative mechanisms. It is also possible that the statistical power of eQTL studies may limit their ability to identify these genes as true eGenes. Further investigation is warranted to comprehensively understand the regulatory landscape of non-PhyeQTL and their implications for gene expression regulation.

### FoldRec-Based Method for Enhancer Target Prediction

The application of FoldRec-predicted 3D chromatin interactions to identify regulatory eQTL holds promise for extending our understanding of enhancer-gene relationships beyond eQTL loci. The widely adopted activity-by-contact (ABC) model [16] for predicting enhancer-gene connections originally utilize the Hi-C data. However, their subsequent results indicate that alternative approaches of estimating 3D contact based on genomic distance or average values of Hi-C data yield comparable results in predicting CRISPR data. This unexpected outcome underscores the need for a critical reassessment of how Hi-C data is utilized, suggesting that current methodologies may not fully exploit the rich insights offered by 3D chromatin interaction studies.

Distinguishing the presence or absence of physical interactions between enhancers and genes from Hi-C data poses a significant challenge. Rather than relying solely on all Hi-C interactions, we leverage FoldRec interactions to predict enhancer-gene interactions. Evaluation on CRISPR-based experimental data provided by Fulco et al. [16] demonstrates the superior performance of our FoldRec-based method in predicting long-range enhancer-gene connections compared to the ABC model (see Fig. 8). These results highlight the potential of FoldRec interactions in predicting functional enhancer-gene relationships, providing a promising avenue for advancing our understanding of gene regulation within the framework of chromatin architecture.

**Figure 8:**
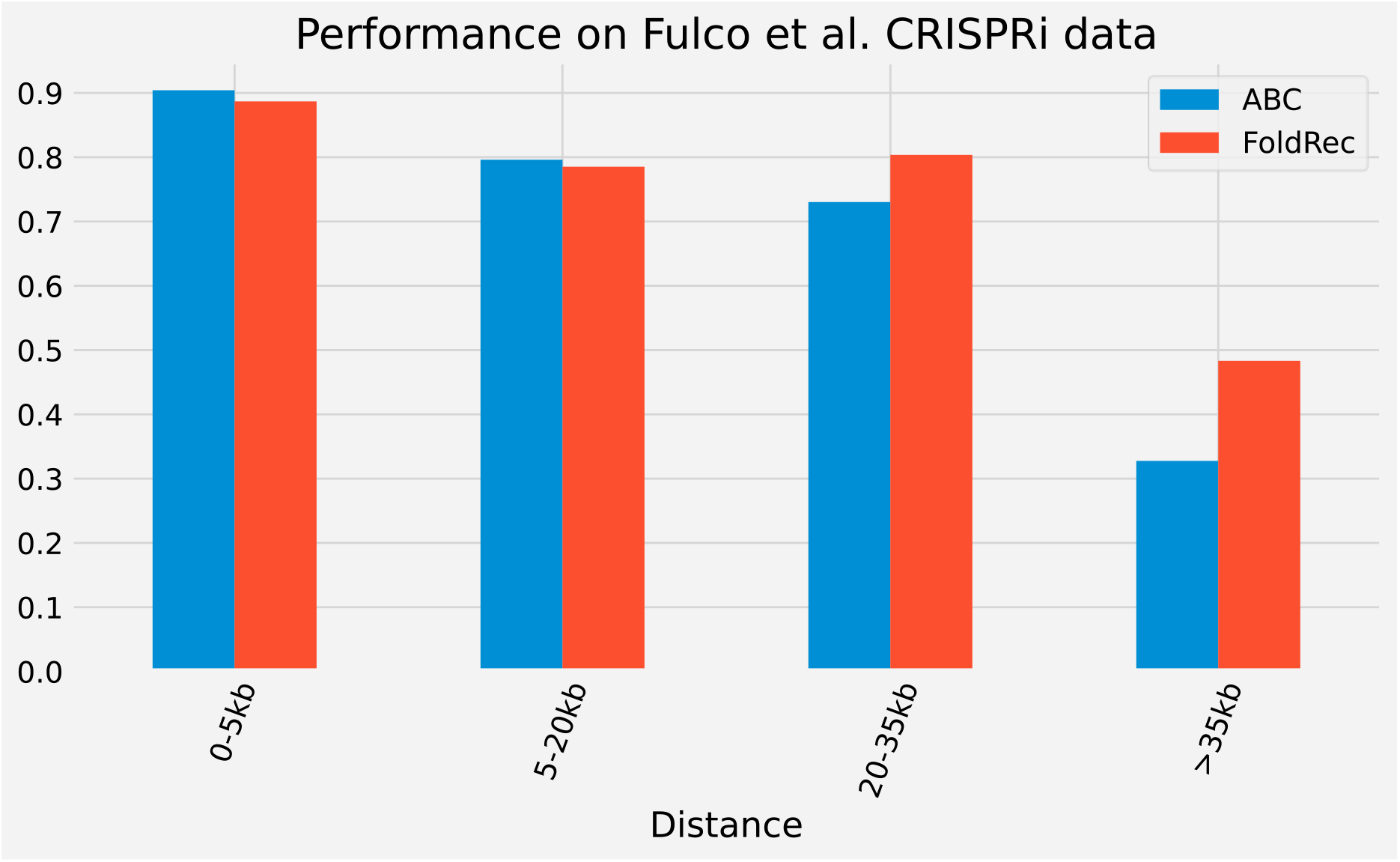
Comparison of FoldRec enhancer target prediction with ABC model. Enhancer–gene pair classification performance (CRISPRi-validated versus non validated candidate enhancers), stratified by relative distance between enhancer–gene pairs. Our model performs better particularly in predicting long-range targets of enhancers.

## Discussion

We have introduced a novel computational framework that integrates genetic (eQTL) and chromatin interactions (Hi-C) to establish physical and functional connections between genetic variants and their target genes. Although both Hi-C and eQTL methodologies provide complementary insights into the associations between regulatory elements (CREs) and target genes, it is challenging to distinguish functionally significant interactions from the vast number of interactions obtained in a Hi-C data.

Our method, based on FoldRec interactions, selectively identifies a small subset of significant Hi-C interactions. These interactions are then overlapped with eQTL-eGene interactions to identify a subset of eQTLs variants that exhibit both genetic and physical interactions, referred as PhyeQTLs. They exhibit enrichment for both statistically and experimentally identified causal variants. Additionally, PhyeQTLs have strong regulatory characteristics, including enrichmnet with open chromatin and tissue-specific transcription factor binding sites, and are associated with tissue-specific PhyeGenes. These findings show that we propose PhyeQTLs can be used as a physical proximity-based method for fine-mapping functional eQTLs.

Validation through CRISPR-based experiments confirms the target genes identified by PhyeQTLs, underscoring their functional significance over other eQTLs. Furthermore, FoldRec interactions also can uncover putative target genes of non-PhyeQTLs which are beyond their annotated eGenes, validated through CRISPR-based experiments. The efficacy of enhancer target prediction based on FoldRec interactions for long range interactions further demonstrates the utility of FoldRec interactions beyond traditional eQTL loci.

Despite the strengths of our approach, certain limitations exist. We interchangeably utilize primary tissue-based data and matching cell line-based data, potentially affecting the degree of reliability of our results. Additionally, a resolution gap exists between eQTL-eGene interactions and Hi-C interactions, which could be addressed with the availability of high-resolution Hi-C data to improves the accuracy of our findings.

## Methods

### Data description

We obtained eQTL data from the GTEx Analysis V8 release using the ‘signif_variant_gene_pairs.txt.gz’ files for four tissues: Cells-EBV-transformed lymphocytes (LCL), Whole Blood (WHLBLD), Lung, and Liver. Hi-C datasets were downloaded from the 4DN Data Portal for the cell lines GM12878, K562, IMR90, and HepG2. Additionally, we downloaded ATAC-seq, RNA-seq, ChIP-seq for H3K27ac, and ChromHMM annotation data from the ENCODE Data Portal. Detailed dataset information is provided in Supplementary Table 1. We analyzed eQTL data for the four GTEx tissues and used matching cell line data when tissue-specific data were unavailable.

### Genomic assembly

All data is based on GRCh38/hg38 genome assemblies unless otherwise specified.

## FoldRec interactions identification

### Constructing a physical null model of chromatin chains

We model random chromatin fibers as self-avoiding polymer chains consisting of beads, employing a fractal Monte Carlo sampling approach. Specifically, we generated a random ensemble of 300,000 polymer chains, sampled uniformly from all geometrically feasible chains confined within the nuclear volume. Each polymer chain comprises 400 spherical monomer beads. These beads, each with a diameter of approximately 40 nm, represent ∼5 kb of DNA, matching the resolution of Hi-C data. To account for confinement and volume exclusion effects, each self-avoiding polymer chain was constrained to reside within a spherical nuclear volume specific to the corresponding cell type. For GM12878, the nuclear volume was taken as 237 μm^3^; for K562, 904 μm^3^; for IMR90, 381 μm^3^; and for HepG2, 850 μm^3^. The nuclear volume of each cell type was then scaled to preserve a constant base pair density, resulting in volumes of 608 nm^3^, 822 nm^3^, 616 nm^3^, and 805 nm^3^ for GM12878, K562, IMR90, and HepG2, respectively. The contact frequency matrix of the randomly sampled chromatin chains was obtained by counting the frequency of interacting monomer pairs. Two chromatin monomers were considered interacting if their Euclidean distance was ≤80 nm.

### Calculating *p*-values for calling FoldRec interactions

To determine the statistical significance of each Hi-C measured interaction, we employ a scalable Bag of Little Bootstraps resampling procedure over the uniform random 3-D polymer ensemble, using 10,000 outer replicates to obtain a null distribution of random chromatin contacts. For each cell type, p-values are assigned to each Hi-C contact frequency based on the proportion of bootstrap replicate contact frequencies that exceed the measured Hi-C contact frequency at the same genomic distance for the corresponding cell type. Finally, to control for multiple testing, a Hi-C interaction is deemed significant (called FoldRec) if the FDR-adjusted p-value is less than 0.05.

## Code availability

Source code for null model chromatin folding by fractal Monte Carlo is available via git repository at https://bitbucket.org/aperezrathke/chr-folder.

## Acknowledgements

This work is supported by NIH grants 1R03OD032628-01, 1R03OD036492-01 and 2R35GM127084-06. An award for computer time was provided by the U.S. Department of Energy’s (DOE) Innovative and Novel Computational Impact on Theory and Experiment (INCITE) Program. This research used resources from the Argonne Leadership Computing Facility, a U.S. DOE Office of Sciensce user facility at Argonne National Laboratory, which is supported by the Office of Science of the U.S. DOE under Contract No. DE-AC02-06CH11357. We thank the labs in the 4DN and ENCODE consortium for generating the data.

