## Supplementary Figures for "A New Approach for Discovering Functional Links Connecting Non-Coding Regulatory Variants to Gene Targets"

**Figure S1**

**A** GTEx Tissues and their macthed cell lines for Hi-C data

| Tissues | Cell Lines |
| --- | --- |
| Cells - EBV-transformed lymphocytes (LCL) | GM12878 |
| Whole Blood (WHLBLD) | K562 |
| Lung | IMR90 |
| Liver | HepG2 |

**B**

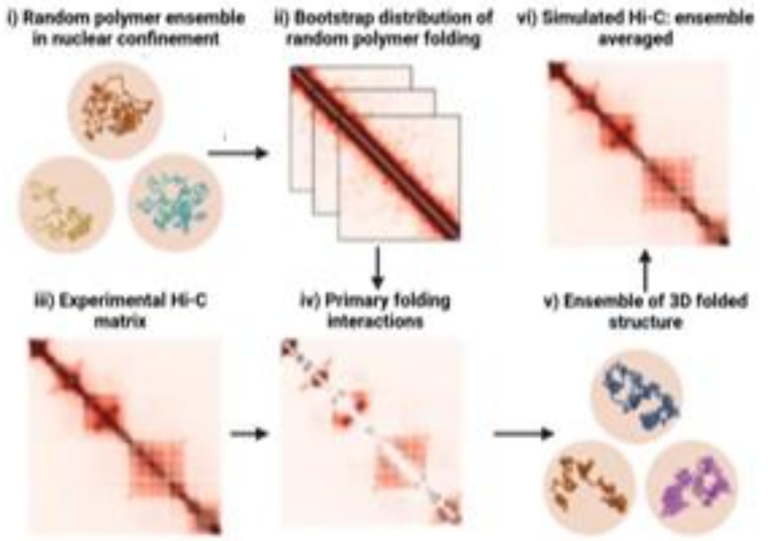


**C**

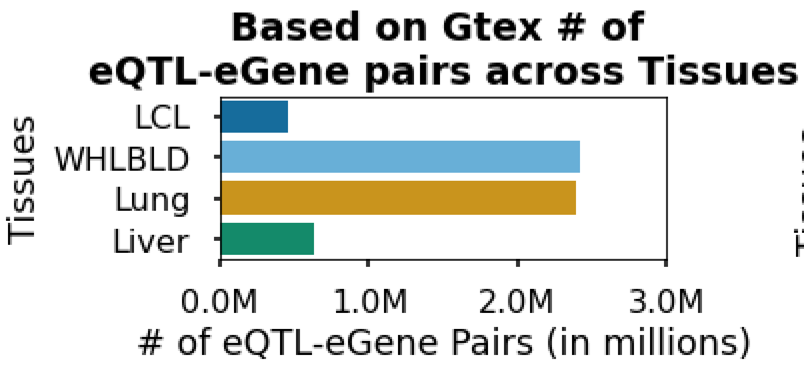


**D**

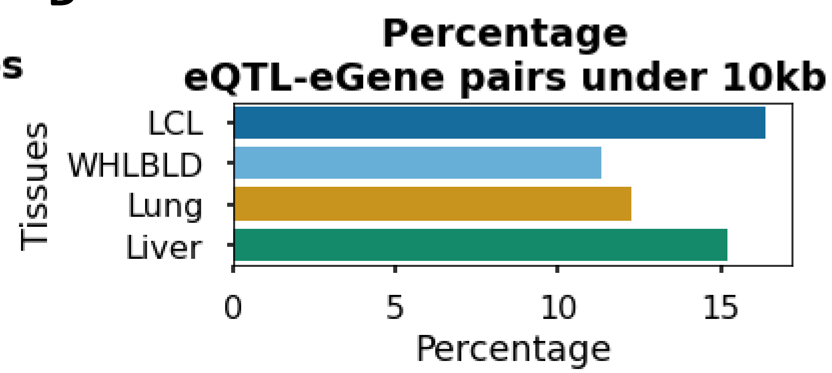


**E**

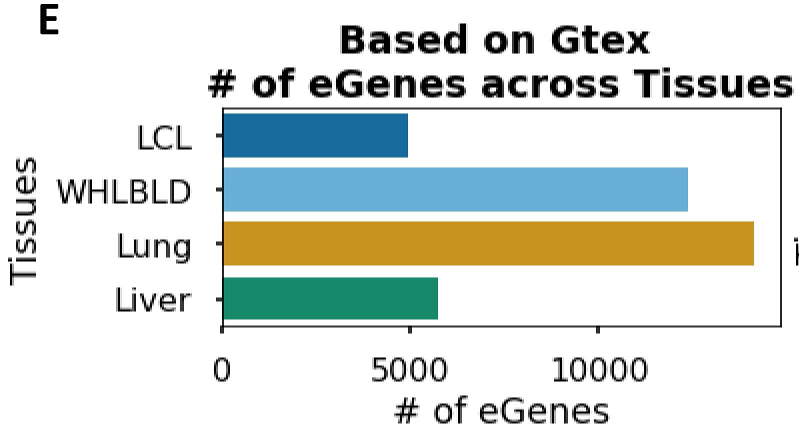


**F**

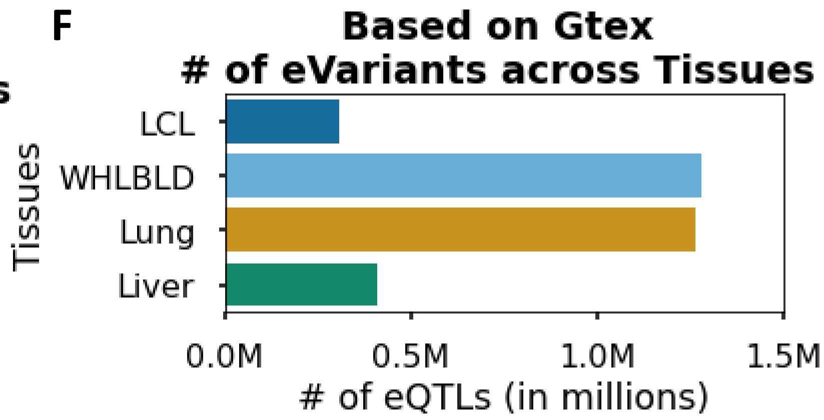


**G**

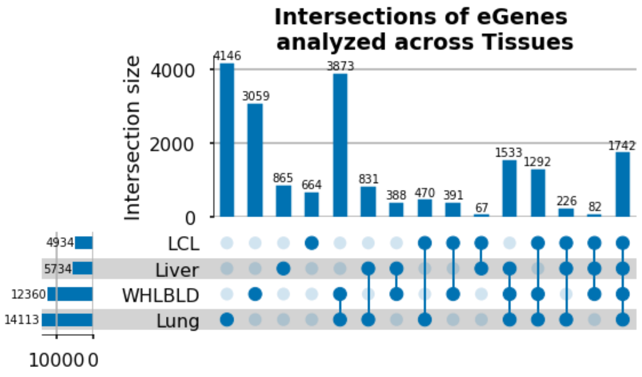


**H**

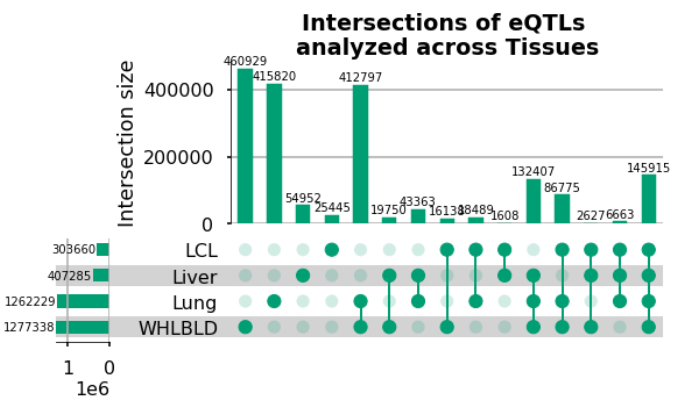

### Figure S2

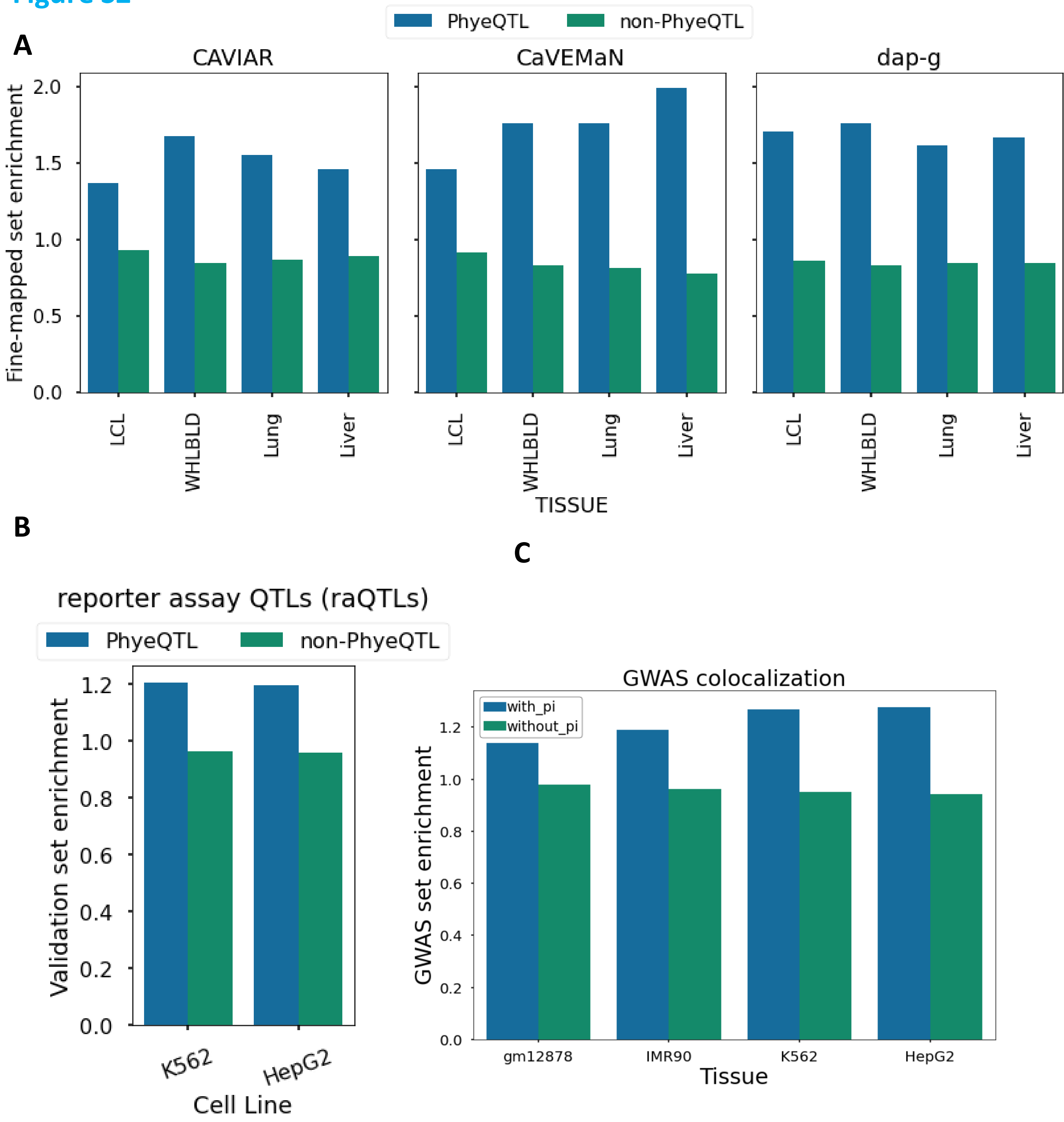
A
PhyeQTL
non-PhyeQTL
CAVIAR
CaVEMaN
dap-g
Fine-mapped set enrichment
0.0
0.5
1.0
1.5
2.0
LCL
WHLBLD
Lung
Liver
TISSUE
B
reporter assay QTLs (raQTLs)
Validation set enrichment
0.0
0.2
0.4
0.6
0.8
1.0
1.2
K562
HepG2
Cell Line
C
GWAS colocalization
with_pi
without_pi
GWAS set enrichment
gm12878
IMR90
K562
HepG2
Tissue

**Figure S3**

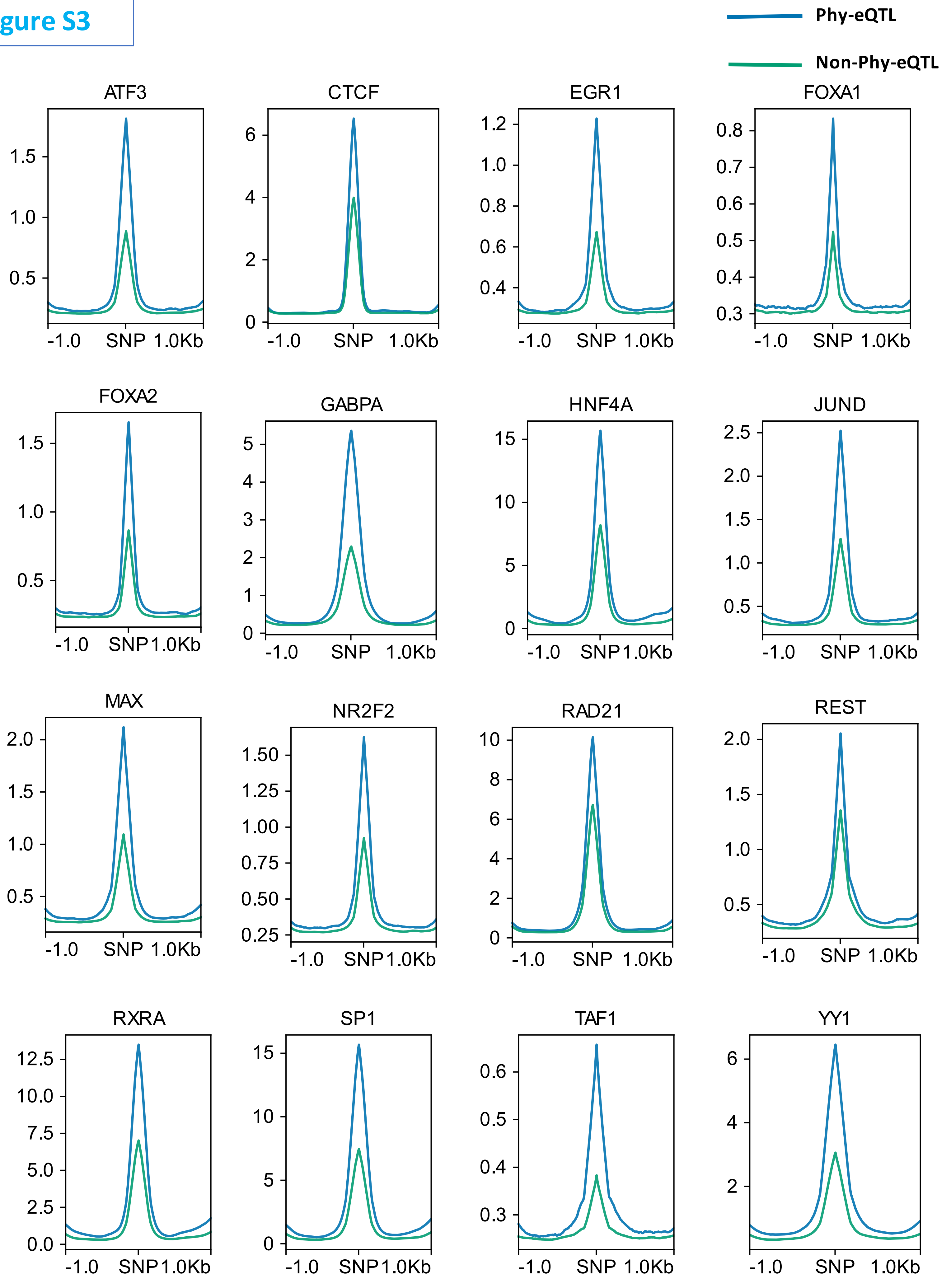
Phy-eQTL
Non-Phy-eQTL
ATF3
CTCF
EGR1
FOXA1
FOXA2
GABPA
HNF4A
JUND
MAX
NR2F2
RAD21
REST
RXRA
SP1
TAF1
YY1
-1.0
SNP
1.0Kb

### Figure S4

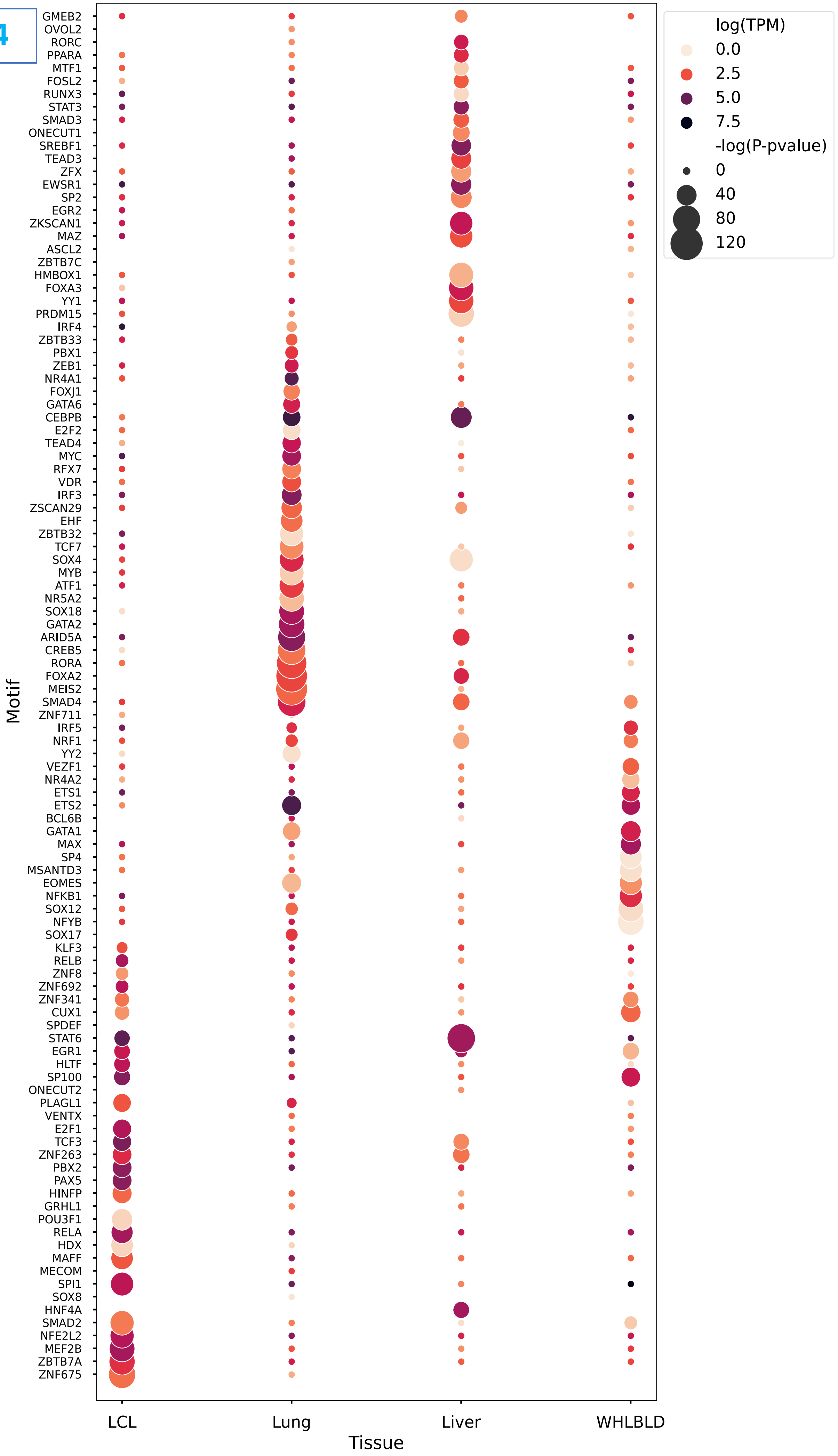

log(TPM)
0.0
2.5
5.0
7.5
-log(P-pvalue)
0
40
80
120
Motif
GMEB2
OVOL2
RORC
PPARA
MTF1
FOSL2
RUNX3
STAT3
SMAD3
ONECUT1
SREBF1
TEAD3
ZFX
EWSR1
SP2
EGR2
ZKSCAN1
MAZ
ASCL2
ZBTB7C
HMBOX1
FOXA3
YY1
PRDM15
IRF4
ZBTB33
PBX1
ZEB1
NR4A1
FOXJ1
GATA6
CEBPB
E2F2
TEAD4
MYC
RFX7
VDR
IRF3
ZSCAN29
EHF
ZBTB32
TCF7
SOX4
MYB
ATF1
NR5A2
SOX18
GATA2
ARID5A
CREB5
RORA
FOXA2
MEIS2
SMAD4
ZNF711
IRF5
NRF1
YY2
VEZF1
NR4A2
ETS1
ETS2
BCL6B
GATA1
MAX
SP4
MSANTD3
EOMES
NFKB1
SOX12
NFYB
SOX17
KLF3
RELB
ZNF8
ZNF692
ZNF341
CUX1
SPDEF
STAT6
EGR1
HLTF
SP100
ONECUT2
PLAGL1
VENTX
E2F1
TCF3
ZNF263
PBX2
PAX5
HINFP
GRHL1
POU3F1
RELA
HDX
MAFF
MECOM
SPI1
SOX8
HNF4A
SMAD2
NFE2L2
MEF2B
ZBTB7A
ZNF675
LCL
Lung
Liver
WHLBLD
Tissue

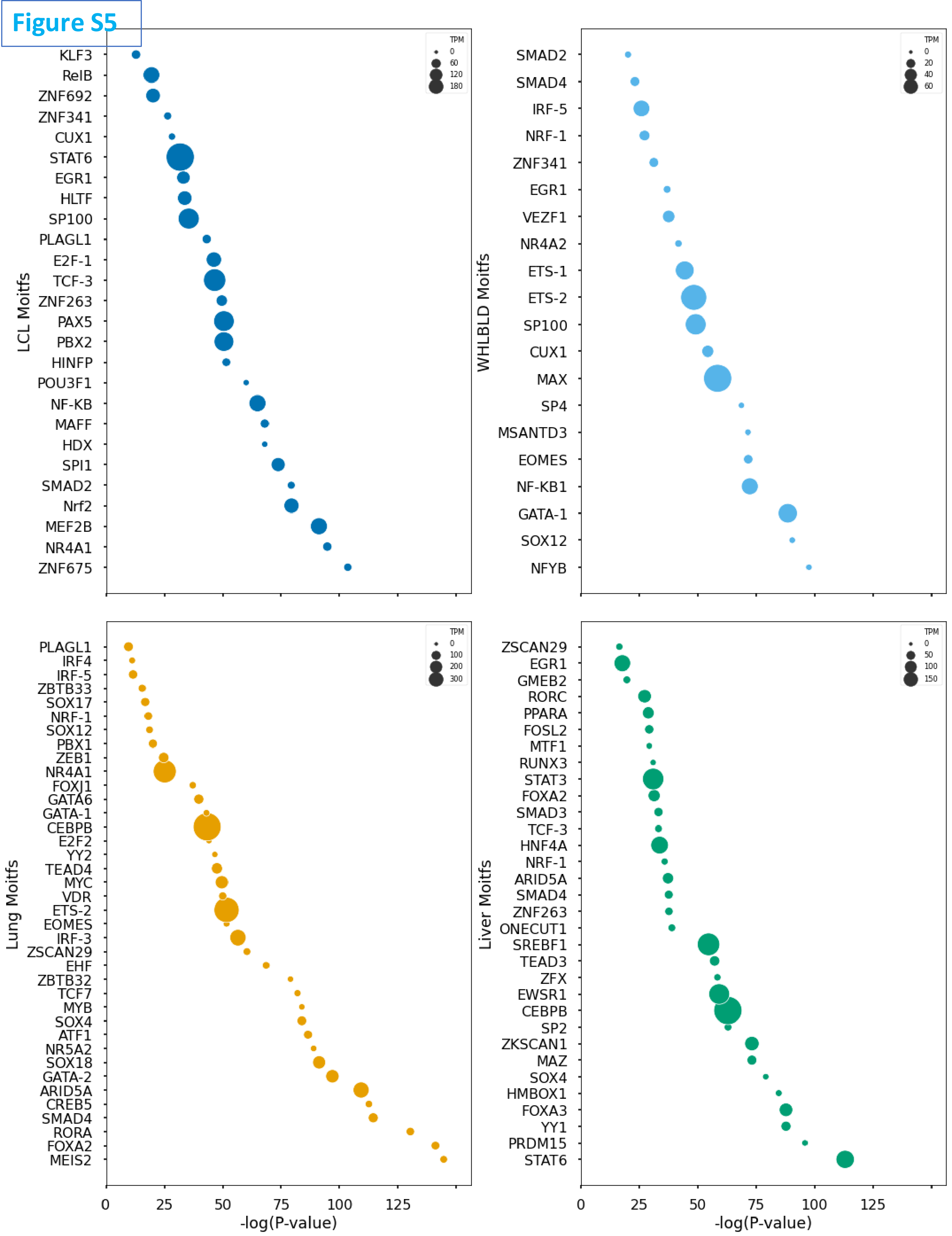
Figure S5
LCL Moitfs
KLF3
RelB
ZNF692
ZNF341
CUX1
STAT6
EGR1
HLTF
SP100
PLAGL1
E2F-1
TCF-3
ZNF263
PAX5
PBX2
HINFP
POU3F1
NF-KB
MAFF
HDX
SPI1
SMAD2
Nrf2
MEF2B
NR4A1
ZNF675
TPM
0
60
120
180
WHLBLD Moitfs
SMAD2
SMAD4
IRF-5
NRF-1
ZNF341
EGR1
VEZF1
NR4A2
ETS-1
ETS-2
SP100
CUX1
MAX
SP4
MSANTD3
EOMES
NF-KB1
GATA-1
SOX12
NFYB
TPM
0
20
40
60
Lung Moitfs
PLAGL1
IRF4
IRF-5
ZBTB33
SOX17
NRF-1
SOX12
PBX1
ZEB1
NR4A1
FOXJ1
GATA6
GATA-1
CEBPB
E2F2
YY2
TEAD4
MYC
VDR
ETS-2
EOMES
IRF-3
ZSCAN29
EHF
ZBTB32
TCF7
MYB
SOX4
ATF1
NR5A2
SOX18
GATA-2
ARID5A
CREB5
SMAD4
RORA
FOXA2
MEIS2
TPM
0
100
200
300
0
25
50
75
100
125
-log(P-value)
Liver Moitfs
ZSCAN29
EGR1
GMEB2
RORC
PPARA
FOSL2
MTF1
RUNX3
STAT3
FOXA2
SMAD3
TCF-3
HNF4A
NRF-1
ARID5A
SMAD4
ZNF263
ONECUT1
SREBF1
TEAD3
ZFX
EWSR1
CEBPB
SP2
ZKSCAN1
MAZ
SOX4
HMBOX1
FOXA3
YY1
PRDM15
STAT6
TPM
0
50
100
150
0
25
50
75
100
125
-log(P-value)
